# Neutrophils degranulate GAG-containing proteoglycofili, which block *Shigella* growth and degrade virulence factors

**DOI:** 10.1101/2022.02.08.479570

**Authors:** Antonin C André, Marina Valente Barroso, Jurate Skerniskyte, Mélina Siegwald, Vanessa Paul, Lorine Debande, Caroline Ridley, Isabelle Svahn, Stéphane Rigaud, Jean-Yves Tinevez, Romain R Vivès, Philippe J Sansonetti, David J Thornton, Benoit S Marteyn

## Abstract

Neutrophil degranulation plays a central role in their ability to kill pathogens but also to stimulate other immune cells^1–3^. Here we show that neutrophil degranulation, induced in hypoxia or upon *Shigella* infection *in vitro* and *in vivo*, leads to the release of polymers called neutrophil Proteoglycofili (PGF). PGF are mainly composed of granular proteins (myeloperoxidase, elastase, lactoferrin, cathelicidin, albumin) pre-stored in various types of granules, and chondroitin sulfate. PGF individual fibers have a diameter of 43.9 ± 20.3 nm and. They secreted by viable neutrophils and do not contain DNA, as opposed to NETs which contains also granular proteins, chondroitin sulfate in addition to chromatin, released upon neutrophil disintegration and cell death.

We demonstrated that PGF block the growth of *Shigella* and other bacteria and degrade *Shigella* virulence factors. The degradation of the chondroitin sulfate polymers with testes hyaluronidases destabilizes PGF ultrastructure and abolishes its antimicrobial activity. Our results provide novel insights in the neutrophil degranulation process and open new doors for the investigation of PGF contribution to cytokines concentration gradient formation and adaptive immune cells activation. Further investigations are required to better appreciate the importance of this “sterile blaster” in infectious or inflammatory diseases.

## Results

Neutrophil degranulation is induced by the mobilization of azurophil (α), specific (β1), gelatinase (β2) granules and secretory vesicles (γ). It was long considered that the mobilization of different types of granules was differential and that degranulated proteins diffused in the extracellular microenvironment.

Neutrophils are mainly glycolytic cells^4^ and are well-adapted to low-oxygen microenvironments^5^ We previously demonstrated that neutrophil viability was maintained when cells were purified and cultured under anoxia^6,7^, allowing the study of their physiology *in vitro* during extended periods of time. Applying this new method (Fig. 1a), we observed that a dense oligomeric material composed of polysaccharides or glycoproteins was released by neutrophils under anoxia, as revealed by blue alcian and periodic acid-Schiff (PAS) staining (Fig. 1b). According to its composition and ultrastructure, this extracellular network was named neutrophil proteoglycofili (PGF). The integrity of PGF, like other mucoidal biomaterials, was only preserved upon fixation with a Carnoy’s solution, not with alternative fixative solutions, such as paraformaldehyde (Extended Data Fig. 1a). By SEM, we determined that the mean diameter of individual polymers composing PGF was 43.9 ± 20.3 nm (Fig. 1c). We confirmed in a kinetic study, that neutrophils secreting PGF under anoxic conditions remained viable (Extended Data Fig. 1b, until t=18h, *p*>0.5). Interestingly, neutrophil PGF were labeled with the Myelotracker peptide, a specific marker of lactoferrin^8^, both *in vitro* (in neutrophil cultures, Fig. 1d). Under basal conditions, lactoferrin is stored in neutrophil secondary (β1) and specific granules (β2): its detection within PGF strongly suggests that neutrophil degranulation sustains PGF formation. These results are consistent with previous reports showing that neutrophil degranulation is promoted under low-oxygen conditions^9^. In addition, PGF were detected *in vivo* within hypoxic foci of infection formed by the pathogenic enterobacteria *Shigella* within the guinea pig colonic mucosa^10^ (Fig. 1e and Extended Data Fig. 1d).

**Figure 1.**
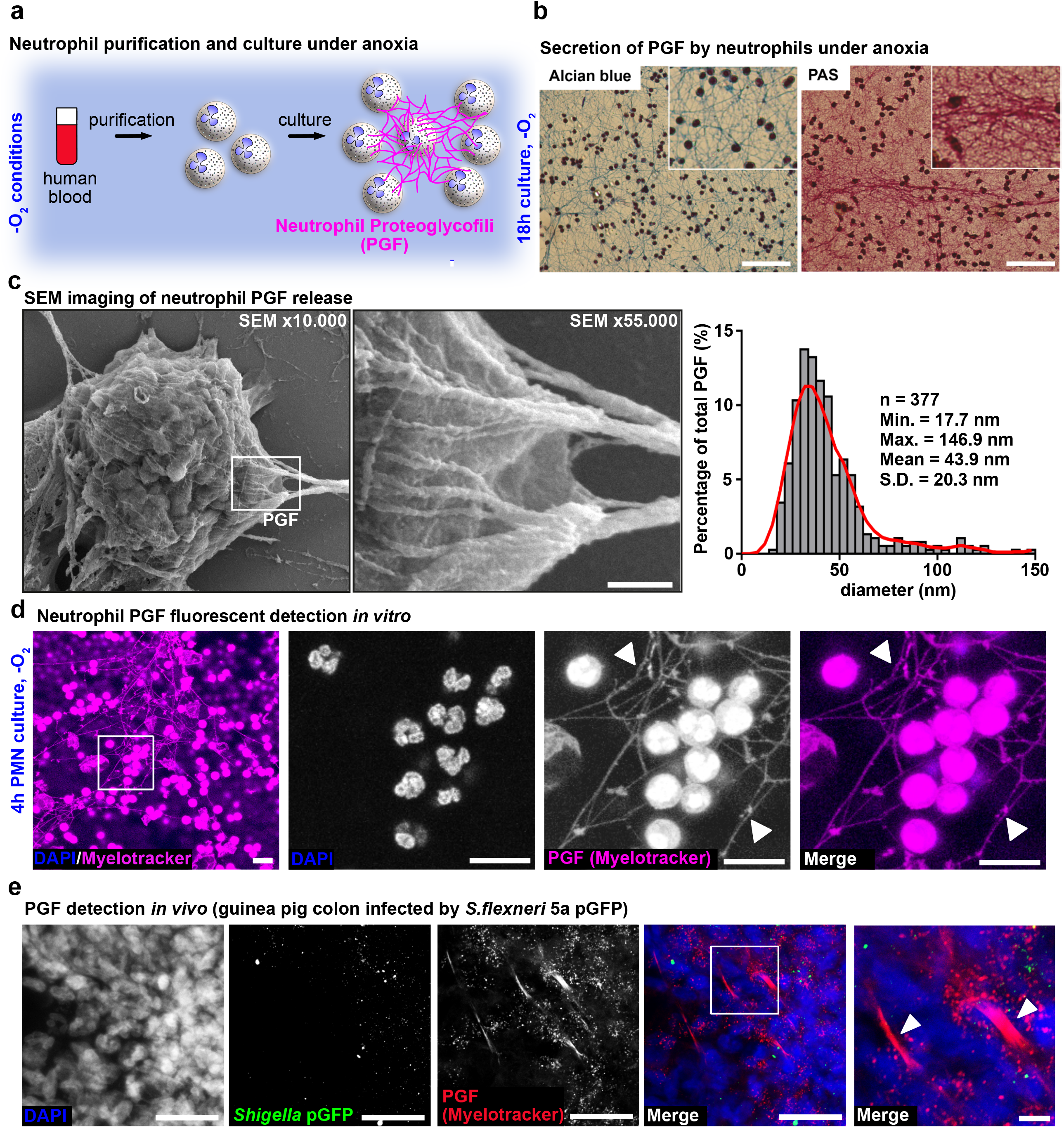
Neutrophil extracellular proteoglycofili (PGF) are released by viable cells and contain lactoferrin. **(a)** PGF are released by neutrophils purified and cultured for 18h, in an optimized culture medium (RPMI 1640 + 10 mM Hepes + 3 mM glucose), under anoxic conditions to maintain their viability ^6^. **(b)** PGF are composed of neutrophil secreted glycoproteins, as revealed upon fixation in Carnoy’s solution and staining with PAS and Alcian Blue (18h culture, -O_2_). Bars, 100 µm. **(c)** Neutrophils secreting PGF were fixed and imaged by scanning electron microscopy (SEM). Bar, 300 nm. (See additional PGF and NETs SEM imaging in Extended Data Fig. 3a.). PGF diameter was measured by quantitative image analysis, using the Fiji Software (n=377). **(d)** Neutrophils secreting PGF (4h culture, -O_2_). were fixed in a Carnoy’s solution and stained with Myelotracker-Cy5 (1 µg/mL, magenta), a specific lactoferrin marker^8^, and DNA was stained with DAPI (blue). Bars, 40 µm. **(e)** PGF was detected *in vivo* with Myelotracker-Cy3 (red) within the guinea pig colonic mucosa infected by *Shigell*a pGFP (green). DNA was stained with DAPI (blue). Bars, 40 µm

Our aim was to define the PGF biochemical composition and to investigate its structural organization and function. PGF were purified from neutrophils in anoxic cultures by differential centrifugations. The oligomeric organization of purified PGF was confirmed by TEM (Fig. 2a) and we demonstrated by mass spectrometry that PGF were mainly composed of neutrophil granular proteins (Extended Data Fig. 2a). The most abundant PGF proteins were myeloperoxidase, elastase, lactoferrin, cathelicidin and albumin (Fig. 2b): their presence within PGF was confirmed by western blot analysis (using SDS-Agarose gel instead of SDS-Page gel electrophoresis for quantitative detection, due to the PGF high molecular-weight, Extended Data Fig. 2b) and by immunofluorescence imaging *in vitro* (Fig. 2c). PGF proteins were found to be accumulated within *Shigella* infectious sites *in vivo* (Extended Data Fig. 2c). No modulation of the expression of the corresponding genes was observed in neutrophils secreting PGF (RNAseq analysis, Extended Data, Fig. 2d). These results demonstrated that all types of neutrophil granules were mobilized during PGF formation, regardless to the hierarchy in mobilization among granules^11^.

**Figure 2.**
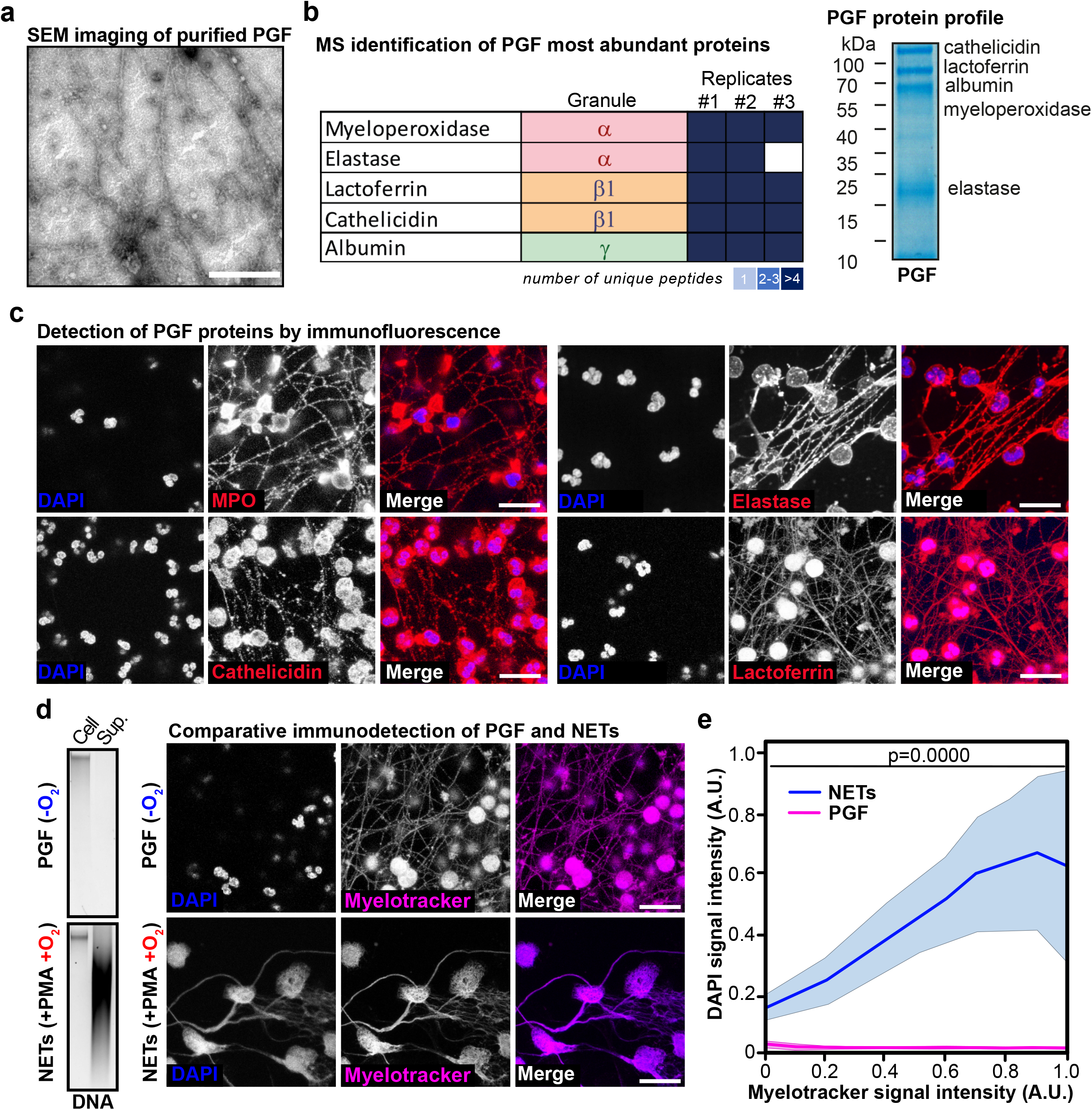
PGF are composed of neutrophil granular proteins. **(a)** PGF was purified from neutrophil cultures by differential centrifugation. PGF ultrastructure was determined by SEM. Bar, 400 nm. **(b)** The most abundant proteins in PGF are lactoferrin, myeloperoxidase (MPO), elastase, cathelicidin (LL-37) and albumin, as determined by MS (n=3, see also Extended Data Fig. 2a-c). The sub-cellular localization of each protein is indicated (α, β1, β2 or γ granules). A representative PGF protein profile is shown on the left-hand panel (4-12% SDS Page gel, Coomassie staining). **(c)** MPO, cathelicidin, lactoferrin and elastase were immunodetected with specific antibodies within PGF secreted by neutrophils (4h culture, -O_2_, Carnoy’s fixation). This result was confirmed by western blot (see Extended Data Fig. 2b) and *in vivo* upon *Shigella* infection (see Extended Data 2c). DNA was stained with DAPI (blue). Bars, 30 µm. **(d-e)** PGF differs from NETs. PGF do not contain DNA, as shown by detecting extracellular DNA in neutrophil cultures (supernatant, sup.) upon PGF secretion (4h, -O_2_) as compared to NETs release (4h, +O_2_, +PMA). This result was confirmed by immunofluorescence imaging and quantitative image analysis. In both conditions, neutrophil cultures were fixed in a Carnoy’s solution and stained with Myelotracker-Cy5 (magenta) and DAPI (blue). Bars, 40 µm. Only extracellular NETs were co-stained with DAPI and Myelotracker-Cy5, as opposed to PGF (p=0.0000).

Neutrophil PGF are released by viable neutrophils (Extended Data Fig. 1b), as opposed to Neutrophil Extracellular Traps (NETs) formation, which is associated with cell death, cell disintegration and the release of the nuclear chromatine^12^. The cell integrity of neutrophils secreting PGF was confirmed by SEM (Extended Data Fig. 3a). We demonstrated by immunofluorescence imaging that neutrophils secreting PGF have multilobulated shape nuclei (Fig. 2d). As opposed to PMA-induced NETs, no extracellular DNA was detected within PGF (Fig. 2d). This result was confirmed by quantitative imaging analysis, co-staining DNA with Dapi and PGF with Myelotracker (Fig. 2e, p=0.0000). PGF were consistently not susceptible to DNase I but were degraded by Proteinase K (Extended Data Fig. 3b).

To investigate the PGF antimicrobial activity we studied its impact on the virulence and the growth of *Shigella*. The invasion of the epithelium by *Shigella* is mediated by an oxygen-dependent Type 3 secretion system (T3SS)^13^. *Shigella* exports additional virulence factors belonging to the Type 5 secretion system (T5SS) (SepA, Pic and IcsA) to colonize and disseminate within infected tissues^14,15^. Accumulation of PGF was observed *in vivo* within *Shigella* infectious foci (Fig. 1e) and we demonstrated that PGF release was consistently induced in neutrophils infected by *Shigella in vitro*, regardless of the oxygen availability (Fig. 3a and Extended Data Fig. 4a). In more details, the interaction between *Shigella* and PGF was confirmed by 3D-reconstitution imaging, using the Imaris software (Fig. 3b)

**Fig. 3.**
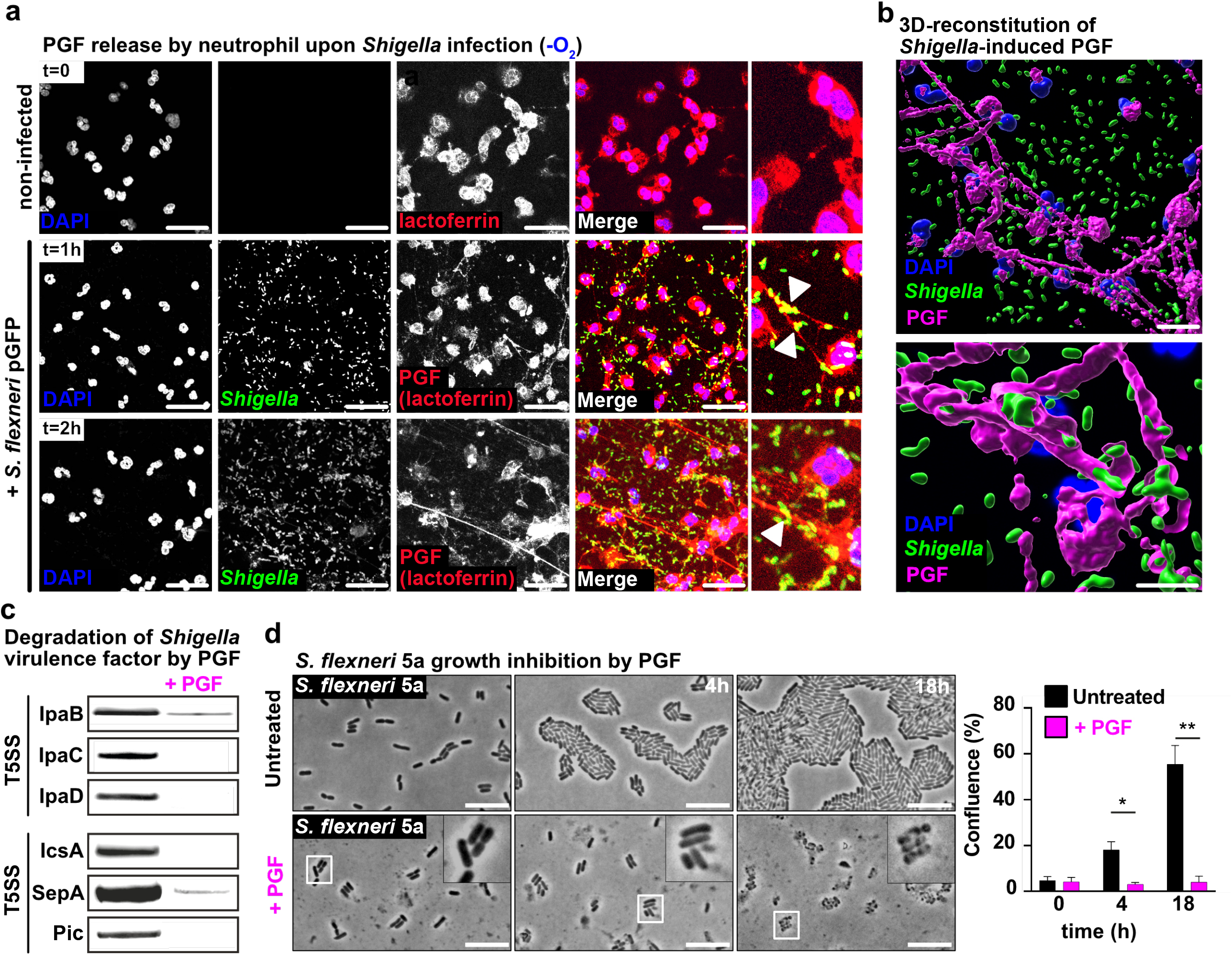
*Shigella* infection stimulates the secretion of neutrophil PGF, which bind to bacteria, degrade virulence factors and block bacterial growth. **(a)** *Shigella* infection induces neutrophil PGF secretion in -O_2_ conditions, as revealed by immunostaining. Similar results were obtained in +O_2_ conditions (see Extended Data Fig. 4a). *Shigella flexneri* 5a was labelled with a specific α-LPS antibody (green), PGF was stained with Myelotracker-Cy3 (red) and DNA was stained with DAPI. Bars, 50 µm. **(b)** 3D segmentation of DAPI, *Shigella*, and PGF using the Surface tool of Imaris 9.6 (BitPlane), and a blow-up with an oblique slicer on the PGF surface to observe *Shigella* located within PGF filaments. Bars, 30 µm (top) and 10 µm (bottom). **(c)** The degradation by PGF of *Shigella* virulence factors (IpaB, IpaC, IpaD, IcsA, SepA) or *Shigella flexneri* 2a (Pic) were observed after 1h incubation at 37°C, as revealed by Coomassie staining (SepA and Pic) or by western blot (IpaB, IpaC, IpaD and IcsA) **(d)** *S. flexneri* 5a growth inhibition by PGF was demonstrated by culturing bacteria on agar pads in the presence of PGF (37°C, 4h and 18h) Bar, 5µm. The confluence of bacterial cultures was quantified using the Fiji software (n=3, * indicates *p* < 0.05 and ** indicates *p* < 0.01(Student’s T-test).

We demonstrated that PGF degrade *Shigella* virulence factors secreted through the T3SS (IpaB, IpaC and IpaD) or belonging to the T5SS (IcsA, SepA and Pic) (Fig. 3c). PGF proteolytic activity is inhibited in the presence of a protease inhibitor cocktail or upon heat inactivation (Extended Data Fig. 4b). It has been previously reported that *Shigella* virulence factors were degraded by elastase^3^, which may be responsible of this PGF activity. We confirmed that as control, PGF did not degrade important human proteins involved in the host anti-microbial response such as lactoferrin, and the heavy and light chains of IgG and IgA (Extended Data Fig. 4c).

We demonstrated that PGF blocked *Shigella flexneri* 5a growth *in vitro*, in the presence of oxygen (Fig. 3d, 18h *p*<0.01). The integrity of bacteria appeared to be altered by PGF, as observed by phase contrast microcopy. PGF also blocked to the same extent *E. coli* K12, *Shigella flexneri* 2a and *Shigella sonnei* growth (Extended Data Fig. 5a, 18h *p*<0.001). We confirmed that the antimicrobial activity of PGF was maintained in the absence of oxygen, using *Shigella flexneri* 5a as a target (Extended Data Fig. 5b, +O_2_ *p*<0.001 vs -O_2_ *p*<0.01). It must be noticed that PGF elicited adverse effects on human epithelial cells in culture. We demonstrated that when incubated at high yield (300 µg/mL), PGF drastically changed the shape and integrity of epithelial cells *in vitro* (Extended Data Fig. 6a) and decreased cell adhesion and growth rate (Extended Data Fig. 6b, *p*<0.001). As anticipated based on these results, we demonstrated that epithelial cell death was consistently induced by PGF after 6h and 24h incubation (Extended Data Fig. 6c and 6d, *p*<0.01).

We further aimed at identifying PGF components which sustain their fibrillar ultrastructure, which cannot be supported by any protein composing PGF. It has been previously demonstrated in several past studies that neutrophils may produce glycosaminoglycans (GAGs), which are known to form long linear polysaccharides. It was reported using paper electrophoresis that chondroitin sulfate-like substances were detected in neutrophil total cell extracts^16–18^. It was further reported that chondroitin sulfate represents more than 60% of total neutrophil GAGs while heparan sulfate (30%) and hyaluronic acid (10%) represent minor GAGs fractions^19^. We hypothesized that GAGs may be present in PGF and may support their fibrillar ultrastructure. We confirmed this hypothesis by demonstrating using RPIP-HPLC that purified PGF contain chondroitin sulfate, not heparan sulfate (Fig. 4a). This result was confirmed by western blot analysis (Extended Data Fig. 7a). In more details only non-sulfated (non-S) and mono-sulfated (4S/6S) chondroitin were identified in PGF (Fig. 4a). It has to be noticed that chondroitin sulfate (non-sulfated or mono-sulfated) was also identified in NETs (Extended Data Fig. 7b). The secretion of chondroitin sulfate by neutrophils in culture was confirmed *in vitro* by immunofluorescence imaging (Fig. 4b). We could reveal an accumulation of chondroitin sulfate *in vivo*, within *Shigella* foci of infection (Fig. 4c).

**Fig. 4.**
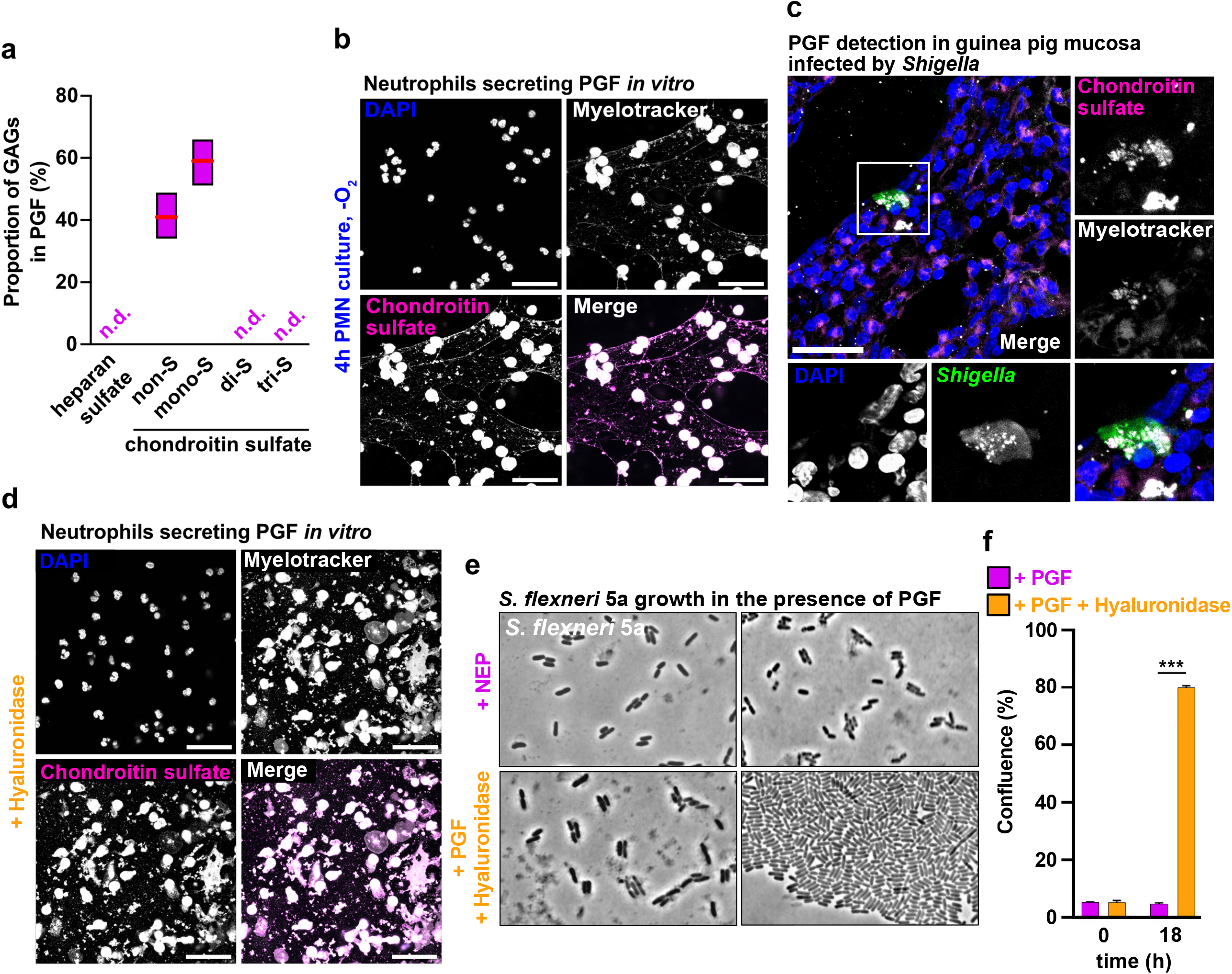
PGF oligomeric structure is sustained by chondroitin sulfate, which is essential for *Shigella* killing. **(a)** The presence of chondroitin/chondroitin sulfate in PGF was confirmed by RPIP-HPLC. Heparan sulfate, chondroitin di- or tri-sulfate were not detected. Results are expressed as percentage of total PGF GAGs (n=3). This result was confirmed by western blot (see Extended Data Fig. 8a) **(b)** Chondroitin sulfate was immunodetected with a specific antibody within PGF secreted by neutrophils (4h culture, -O_2_, Carnoy’s fixation). PGF was stained with Myelotracker-Cy3 (white) and with an anti-chondroitin sulfate antibody (magenta); DNA was stained with DAPI (blue). Bars, 50 µm. **(c)** PGF was immunodetected within the guinea pig colonic mucosa infected by *Shigella* flexneri 5a pGFP (green). PGF was stained with Myelotracker-Cy3 (white) and with an anti-chondroitin sulfate antibody (magenta); DNA was stained with DAPI (blue). Bar, 100 µm. **(d)** PGF secreted by neutrophils was treated with hyaluronidase type I-S. (4h culture, -O_2_, Carnoy’s fixation). PGF was stained with Myelotracker-Cy3 (white) and with an anti-chondroitin sulfate antibody (magenta); DNA was stained with DAPI. Bars, 50 µm. **(e)** *Shigella flexneri* 5a was grown on agar pad with purified PGF treated or not with hyaluronidase type I-S. (37°C, 4h and 18h) Bar, 5µm. **(f)** The confluence of bacterial cultures was quantified using the Fiji software (n=3, * indicates *p* < 0.05 and ** indicates *p* < 0.01(Student’s T-test).

We further aimed at determining the contribution of chondroitin sulfate in PGF structure and antimicrobial functions (degradation of virulence factors and inhibition of bacterial growth). To proceed, PGF was treated with hyaluronidase type I-S from bovine testes, which cleaves β-N-acetylhexosamine-[1→4] glycosidic bonds chondroitin sulfate, chondroitin and hyaluronic acid. We demonstrated that the oligomeric structure of PGF released by neutrophils in culture was altered by a hyaluronidase type I-S treatment (Fig. 4d). In addition, the treatment of purified PGF with this enzyme reverse its capacity to block *S. flexneri* 5a growth (Fig. 4e, *p*<0.001). Taken together, these results confirm the essential role of chondroitin sulfate in PGF structural organization and antimicrobial function.

Together, the data suggest that neutrophil degranulation product defined as PGF is a well-structured compound composed of chondroitin sulfate and granular antimicrobial proteins. PGF is composed of a cocktail of granular proteins stored in all types of granules (α, β1, β2 and γ) suggesting that during PGF release the mobilization of neutrophil granules is not sequential but appeared to be simultaneous. The diffusion of the soluble content of mobilized granules may be limited and is anticipated to accumulate in the close vicinity of activated neutrophils within inflammatory sites. Chondroitin sulfate has a strong affinity for cytokines such as IL-8, CXCL1, CXCL5^20,21^ suggesting that PGF may play a central role in the formation of cytokines concentration gradients *in vivo*, favoring the chemoattraction of immune cells to inflammatory sites. PGF is an additional degranulation compounds which will have to be investigated together with granular proteins to better understand the immunomodulatory role of neutrophil degranulation ^22^.

We observed that PGF and NETs contain granular proteins and chondroitin sulfate at similar yields, although PGF do not contain DNA (Fig. 2d-e) and are released by viable cells, as opposed to NETs formation which is associated with neutrophil cell-death. Further investigations are required to determine whether PGF release is a pre-requisite for NETs formation, considering that NET inducers such as PMA are well-known inducers of neutrophil degranulation^23^. The respective contribution of the chromatin and PGF in NETs structural organization will have to be additionally clarified. In addition, the presence of PGF in NETs raise the question of their contribution to important immunomodulatory functions of NETs, such as the promotion of metastases in cancer through their interaction with tumor cells ^24,25^, the stimulation of macrophage activation including cytokine production in atherosclerosis^26^ or cylic GMP-AMP synthase (cGAS)-dependent stimulation of an interferon response, since elastase has been identified as the main inducer of this pathway ^27^. Finally, the protein composition of activated neutrophil exosomes is similar to PGF (elastase, lactoferrin, myeloperoxidase and MMP-9). Further investigations are required to determine the presence of chondroitin sulfate in these released particles and its potential contribution to their extracellular matrix degradation activity ^28^

Our findings highlight the unrecognized and potentially underappreciated role of GAGs in addition to granular proteins in neutrophil antimicrobial activity or in their capacity to stimulate other immune cells. Our work provides information for understanding the overall function of the neutrophil degranulation and provides original directions for the design of innovative compounds to target neutrophil in inflammatory diseases ^29^.

## Supporting information

Extended informations

