## Extended informations for "Neutrophils degranulate GAG-containing proteoglycofili, which block *Shigella* growth and degrade virulence factors"

**Extended Data**

### Methods

#### Human cell culture

*Blood collection and neutrophil purification.* All participants gave written, informed consent in accordance with the Declaration of Helsinki principles. Peripheral human blood was collected from healthy patients at the ICAReB service of the Pasteur Institute (authorization DC No.2008-68), the Etablissement Français du Sang (EFS) Aquitaine-Limousin (authorization DC No. AC 2013-2025) and from the Etablissement Français du Sang (EFS) de Strasbourg (authorization n°ALC/PIL/DIR/AJR/FO/606). Human blood samples were collected from the antecubital vein into tubes containing sodium citrate (3,8% final) as an anticoagulant.

Blood samples were rapidly transferred into an anoxic chamber (DG250, Don Whitley) for human neutrophil anoxic purification, using oxygen-free media, as described previously[1]. Briefly, whole blood samples were centrifuged at 600 x *g* for 20 minutes. Platelet rich plasma (PRP) was collected and centrifuged at 2650 x *g* for 20 min to form platelet poor plasma (PPP). Blood cells were resuspended in NaCl 0.9% and dextran sulfate (0.72%). After 30 min sedimentation, the neutrophil-containing upper layer was centrifuged at 300 x *g* for 10 min. Pelleted cells were resuspended in 1 mL PPP and were subsequently separated on a 42% Percoll-plasma (GE Healthcare) gradient by centrifugation at 800 x *g* for 20 min. Neutrophils were collected from the pellet with remaining red blood, which were removed using CD235a (glycophorin) microbeads (Miltenyi Biotec). Purified neutrophils were centrifuged at 300 x *g* for 10 minutes. Pelleted cells were resuspended in PPP or RMPI 1640 culture medium supplemented with 10 mM Hepes and 3mM glucose[2], according to experimental settings.

*Epithelial cell culture.* HEp-2 epithelial cells (ref. CCL-23, ATCC) were cultured at 37°C in the presence of 5% $CO_2$ in DMEM medium (ref. 10741574, ThermoFischer Sientific) supplemented with 10 % Fetal Calf Serum (ref. 16010167, ThermoFischer Scientific).

**Bacterial strains and culture**

*Escherichia coli* (K12 MG1655 or HB101) strains were cultured in Lysogeny broth (LB) at 37°C. *Shigella flexneri* 5a (M90T), *Shigella flexneri* 2a and *Shigella sonnei* wild-type strains were grown in Tryptic soy broth (TSB, 211768, ThermoFisher Scientific) medium at 37°C. *Shigella flexneri* 5a pGFP was obtained upon the transformation of the pFpV25.1 plasmid[3].

*E. coli* HB101 strain was transformed with pUC19-SepA (pZK15)[4] (supplied by Dr. Claude Parsot, Institut Pasteur) or pACYC184-Pic (pPic)[5] for the expression and the purification of SepA and Pic (Serine Protease Autotransporters of Enterobacteriaceae or SPATEs).

**Production and purification of PGF and NETs**

*Proteoglycofili (PGF)*. Neutrophils were allowed to secrete PGF upon 4h or 18h culture at 37°C in RPMI1640 medium (ref. 32404-014, ThermoFisher Scientific) supplemented with 10mM Hepes and 3mM glucose (2x$10^6$ cells/mL). When indicated, after 4h culture, DNAseI (ref), proteinase K (ref) or hyaluronidase type I-S from bovine testes (ref H3506, Merck) were added for 1h at 37°C prior fixation in a Carnoy’s solution.

For high yield production of PGF, neutrophils were cultured in 75 $cm^2$ polystyrene culture flasks (Thermofisher Scientific) under anoxic conditions (DG250 workstation, Don Withley). After indicated culture time, neutrophil cultures were collected and centrifuged at 300 x *g* for 5 minutes to pellet cells. PGF were collected from culture supernatant by ultracentrifugation (46 000 x *g* for 50 min). Pelleted PGF were resuspended in 1mL PBS, as previously described for NETs purification[6]. Samples were sonicated before storage at -20°C.

For imaging analysis, neutrophils were allowed to secrete PGF onto glass coverslips in polypropylene 24-well plates (ref. 290-8324-03F, Evergreen Scientific). At each indicated time point, neutrophils were fixed in Carnoy's solution (60% ethanol, 30% chloroform and 10% glacial acetic acid) for 1 hour at room temperature. Sample fixation in PFA 3.3% was tested but did not allow to maintain the structural integrity of PGF. Samples were subsequently washed in PBS prior staining with Alcian blue (ref. B8438, Merck) or Periodic Acid Schiff (PAS) (refs. P7875 and 3952016, Merck). For immunofluorescent labelling, coverslips were carefully washed and permeabilized with PBS-Triton 0.1% and stained with primary antibodies for 1h at room temperature (RT), washed three times in PBS-Triton 0.1%, prior incubation with secondary antibodies for 1h at RT. After three washes in PBS-Triton 0.1%, three washes in PBS and three washes in water, coverslips were mounted with 10µL ProLongGold® (Invitrogen) and allowed to dry overnight at RT. For electron microscopy imaging, fixed samples were rinsed twice in 0.1M phosphate buffer and dehydrated in a series of increasing ethanol concentrations (30 to 100%). Samples were dried via critical point drying (Leica EM CPD300, Austria). Afterwards cover glasses were mounted onto specifics stubs and coated with platinum, using a sputter coater (Q150T, Quorum Technologies, Kent, UK).

*NETs.* Neutrophil Extracellular Traps (NETs) formation was induced in RPMI 1640 medium (supplemented with 10mM Hepes, 3mM Glucose) in the presence of PMA (Phorbol 12-myristate 13-acetate, ref. P1585, Sigma-Amldrich) (0.1µM) during 4 hours at 37°C under atmospheric conditions (21%), as described previously[6,7]. Samples were processed similarly to PGF (for high yield production and for immunofluorescence/electron microscopy imaging).

**Mass spectrometry**

Prior to mass spectrometry samples were prepared via reduction, alkylation, and trypsin digestion, as previously described[8]. After treatment with trypsin, digested peptides were transferred to a 10 kDa MWCO Vivaspin centrifugal concentrating column and centrifuged at 4.500 x *g* until the whole sample volume had been recovered. Digested tryptic peptides were then acidified to pH 2.0 using 1 % (v/v) trifluoroacetic acid, and subsequently purified using C18 ZipTips (Millipore, Durham, UK). Using an UltiMate 3000 Rapid Separation LC coupled to a LTQ Velos Pro mass spectrometer liquid chromatography tandem mass spectrometry (LC-MS/MS) (Thermo Fisher Scientific, Waltham, USA) was performed on peptide containing samples. Firstly, a pre-column of 20 mm x 180 µm was used to concentrate sample peptides. A 75 mm x 250 mm 1.7 µM ethylene bridged hybrid C18 analytical column (Waters, Manchester, UK) was then used to separate peptides in a gradient from 99% solution A (0.1% formic acid in dd$H_2O$) and 1% solution B (0.1% formic acid in acetonitrile) to 25% solution B. Peptides were then automatically selected for fragmentation via a process of data dependant analysis.

The MASCOT search engine was used to analyse the processed mass spectrometry data. Data were searched against the SwissProt database using the following parameters: 1.2 Da mass accuracy for parent ions, 0.6 Da mass accuracy for fragment ions, one missed cleavage site was allowed for, $2^+$ and $3^+$ ions were selected, carbamidomethyl-cysteine was selected as a fixed modification, and methionine oxidation as a variable modification. The data generated from MASCOT was then analysed in a semi-quantitative manner using Scaffold Proteome software (Proteome Software, Portland, USA), allowing for unique peptide counts of identified proteins to be determined.

**Neutrophil glycosaminoglycans (GAG) analysis**

GAG extraction and compositional analysis was performed as described previously[9]. Here, PGF and NETs were digested with 0.2mg/mL proteinase K (Merck) for 1h at 37°C. Potential GAG/protein complexes were dissociated by incubation with 2M NaCl (30 min at room temperature); protein denaturation was achieved by heating samples (100°C for 10 min). After centrifugation, supernatants were recovered, and concentrated to 50-100 µL in 50 mM Tris-HCl pH 7,5 supplemented with 50 mM NaCl and 2 mM $CaCl_2$ (GAG digestion buffer) using a 3kDa cutoff Centricon® (Merck).

GAGs were then exhaustively digested into disaccharides by incubating samples for 48 h at 37°C, with either Chondroitinase ABC (Sigma-Aldrich, 100 mU) for CS/DS analysis, or with a mix of Heparinase I, II and III (Grampian enzymes, 10 mU each) for HS analysis. Compositional analyses were performed by RPIP-HPLC, as previously described[9]. Samples were applied to a C18 reversed phase column equilibrated in a $H_2O$/acetonitrile (8.5%) buffer supplemented with 1.2 mM of ion-pairing tetra-N-butylammonium hydrogen sulphate (TBA), then resolved using a multi-step NaCl gradient calibrated with authentic GAG disaccharide standards (Iduron). On-line post-column disaccharide derivatization was achieved by the addition of 2-cyanoacetamide (0.25%) in NaOH (0.5%), followed by fluorescence detection (excitation 346 nm, emission 410 nm).

**Antibodies and fluorescent dyes**

Lactoferrin was labelled with a rabbit polyclonal antibody (ref. ab77780, Abcam) (1:500 dilution for WB and 1:100 for IF) or with RI-Myelotracker-cy3 or RI-Myelotracker-Cy5, as indicated, used at 1 µg.mL$^{-1}$ [8,10] (see www.myelotracker.com for more technical details). DNA was labelled with 4′,6-Diamidino-2-phenylindole (Dapi, ref. D9542, Merck) and was used at 1 µg.mL$^{-1}$. Actin was labelled with an Alexa Fluor™ 488 Phalloidin (ref. A12379, ThermoFisher Scientific) (1:40 final dilution for IF). Elastase was detected with a rabbit polyclonal antibody (ref. ab68672, Abcam) (1:500 final dilution for WB and 1:100 final dilution for IF). Cathelicidin was detected with a rabbit polyclonal antibody (ref. ab93357, Abcam) (1:1000 final dilution for WB and 1:200 for IF). Myeloperoxidase was detected with a rabbit monoclonal antibody (ref. A0398, Dako) (1:500 final dilution for WB and 1:200 final dilution for IF). Chondroitin sulfate was detected with a mouse monoclonal antibody (ref. C8035, Merck) (1:200 final dilution for WB and 1:100 for IF). Heparan sulfate was detected with a mouse monoclonal antibody (clone F58-10E4, ref: 370255-1, Amsbio) (1:100 final dilution for WB). Keratan sulfate was detected with a mouse monoclonal antibody (clone 373E1, ref: AMS.PRPG-KS-M01, Amsbio) (1:100 final dilution for WB). Murine anti-*Shigella* 5a lipopolysaccharide polyclonal antibody was used for immunofluorescent labelling (1:500 final dilution)[11]. IpaB was detected with a mouse monoclonal antibody (clone H16) kindly supplied by Dr. Armelle Phalipon, Institut Pasteur) (1:1.000 final dilution for WB). IpaC was detected with a mouse monoclonal antibody (clone K24) kindly supplied by Dr. Armelle Phalipon, Institut Pasteur) (1:1.000 final dilution for WB). IpaD and IcsA were detected with rabbit polyclonal antibodies kindly supplied by Dr. Claude Parsot, Institut Pasteur (1:10.000 final dilutions for WB). For immunofluorescence staining, rabbit and mouse secondary antibodies conjugated with an Alexa-568 fluorophore were used (refs. A11036 and A11031, ThermoFisher Scientific) (1:1.000

final dilution). For western-blot analyses, rabbit and mouse secondary antibodies conjugated with horseradish peroxidase were used (refs. 115-035-003 and 111-035-003, Jackson Immuno Research) (1:10.000 final dilution).

**Flow cytometry**

Cell viability was determined by staining with 0.01% propidium iodide (PI, ref. P4170, Marck) in PBS + 2 mM EDTA for 15 min at RT. PI only penetrates into and stains the DNA of non-viable cells. Fluorescence was measured using a FACSCalibur (BD Bioscience) flow cytometer, recording at least 10,000 events. Data were analysed with CellQuest Pro software (BD Biosciences) and were quantified using the FlowJo software (FlowJo, LLC).

**Electrophoresis and western blot**

*SDS-agarose gel.* To separate high molecular-weight protein complexes (as described previously[12]), samples were loaded in 0.8% agarose gel (containing 0.1% SDS). Sample migration occurred at 60V for 3h. The gel was then reduced for 15 min in a SSC (Saline-Sodium Citrate) buffer with 10 mM DTT and washed in a SSC buffer without DTT. Protein transfer onto a PVDF membrane was achieved under vacuum (GE Healthcare™, VacuGene XL Vacuum Blotting Unit) at 50 mbar for 1h30.

*SDS-Page gel.* 4-12% SDS-PAGE gradient gel (NuPAGE; ref NO0322PK2, ThermoFischer Scientific). Migration occurred at 200V for 25 minutes. When indicated, gels were stained with InstantBlue™ Coomassie (ref. ab119211, Abcam) or protein were transferred onto a PVDF membrane (at 350 mA for 180 min) (Mini TransBlot Electrophoretic Transfer Cell, Biorad).

*Agarose gel.* The presence of DNA in cells and in purified PGF and NETs was determined by migrating samples in 1% agarose gel in the presence of propidium iodide, followed by UV revelation (Molecular Imager®, Biorad).

*Western blot.* PVDF membranes were saturated with a solution of Tris buffered saline with tween 20 (TBST) solution (ref. 91414, Merck) containing 5% milk for 1h and incubated overnight with primary antibodies in TBST. Membranes were washed with TBST three times (5 min) and incubated with a secondary antibody for 1h in TBST containing 1% milk. After three additional washes (5 min) antibody binding was detected with chemiluminescence (ECL™ Prime Western Blotting Detection Reagent, GE Healthcare).

**RNAseq analysis**

Neutrophil mRNA were purified from $5.10^7$ neutrophils either freshly purified or cultured for 18h in anoxia in RPMI 1640 medium supplemented with 10 mM Hepes and 3 mM glucose. Cells were washed once with PBS and centrifuged for 5 minutes at 300 x *g*. Pelleted cells were lysed in 350 µL RLT buffer (Qiagen). Lysates were resuspended with a needle to increase the lysis efficiency. TRIzol (3 volumes) was added to samples followed by vortex and short spin. 1/5 of chloroform was added and samples were vortexed and incubated for 3 minutes at room temperature prior transferring into PLG heavy tubes. Samples were centrifuged for 15 min at 12.000 x *g* (4°C) and RNA-containing aqueous solution was transferred into a new tube. RNA were purified using the RNeasy Mini Kit (Qiagen) and quantified with nanodrop and the quality was assessed with Tapestation. RNA samples were analyzed and sequenced by the GENOM'IC platform (Institut Cochin, Paris, France); RNA sequences were analyzed with the Deseq2 method. Results are averaged from three biological triplicates for each condition.

**Guinea pig model of shigellosis**

Young guinea pigs (Hartley, <150g) were previously infected intrarectally with $10^{10}$ CFU exponentially grown *S. flexneri* 5a pGFP to evaluate acute infection (8h p.i)[13]. In this study, ascorbate-deficient guinea pigs were challenged intrarectally with $10^{10}$ CFU exponentially grown *S. flexneri* 5a pGFP and animals were sacrificed 48h p.i.. In this new animal model of shigellosis, infection symptoms are more severe and the colonization of the colonic mucosa by *Shigella* is more prolonged[14] . After animal were sacrificed, infected colons were collected and fixed in 3.3% paraformaldehyde (PFA) for 2 hours. Fixed colon samples were washed in PBS, incubated at 4ºC in PBS containing 15% sucrose for 90 min, followed by incubation in PBS with 30% sucrose overnight. Samples were frozen in OCT (Sakura) on dry ice. 7 μm sections were obtained using a cryostat CM-3050 (Leica). For immunofluorescent labelling, tissues were washed and permeabilized with PBS-Triton 0.1% and stained with primary antibodies in the presence of 1% BSA overnight at 4°C, washed three times in PBS-Triton 0.1%, prior incubation with secondary antibodies for 1h at RT. After three washes in PBS-Triton 0.1%, three washes in PBS and three washes in water, tissues were mounted with 10µL ProLongGold® (Invitrogen) and allowed to dry overnight at RT.

**Neutrophil infection with *Shigella***

Neutrophils were seeded ($2.10^6$ cells/well) onto coverslips in RPMI 1640 medium supplemented with 10 mM Hepes and 3 mM glucose in 24-well plates containing. Cells were infected at a multiplicity of infection (MOI) of 20 with *S. flexneri* 5a pGFP grown until exponentially phase. Bacteria were spun onto cells by centrifugation at 300 x *g* for 10 min. After 1h, 2h and 4h incubation at 37°C, samples were fixed with Carnoy's solution for 2h at room temperature. For immunofluorescent labelling, coverslips were washed and permeabilized with PBS-Triton 0.1% and stained with primary antibodies for 1h at room temperature (RT), washed three times in PBS-Triton 0.1%, prior incubation with secondary antibodies for 1h at RT. After three washes in PBS-Triton 0.1%, three washes in PBS and three washes in water, coverslips were mounted with 10µL ProLongGold® (Invitrogen) and allowed to dry overnight at RT.

**Epithelial cell viability assay**

HEp-2 cells were seeded in 24-well plates onto coverslips at $5.10^5$ cell/mL in 1 mL DMEM supplemented with 10 mM Hepes. After overnight growth, cells were washed with PBS and incubated with purified PGF diluted in a RPMI + 10 mM Hepes medium at two concentrations: low [PGF] corresponds to 30 µg/mL PGF and high [PGF] corresponding to 300 µg/mL PGF. For immunofluorescence imaging, treated cells were washed in PBS and fixed in PFA 3.3%; for flow cytometry analysis, cells were trypsinized and resuspended 300 µL PBS supplemented with 2mM EDTA and propidium iodide.

**Bacterial growth assay**

Overnight bacterial cultures were subcultured in fresh M9 medium (1:100 dilution) at 37°C until a $OD_{600}$ of 0.3 was reached. 1 mL bacterial suspension was centrifuged at 8.500 x *g* for 3 minutes; pelleted bacteria were resuspended in 500 µL PBS. 1 µL bacterial suspension was deposited onto a 1% M9-agar pad and covered with a square coverslip. Agar pads were incubated at 37°C to allow bacterial growth, which was assessed by widefield microscopy at t=0, t=4h and t=18h. When indicated, bacterial growth was performed in an anoxic chamber (-$O_2$ condition). To evaluate the impact of PGF on bacterial growth, when indicated, 5 µg of purified PGF was deposited onto the agar pad prior to depositing the bacterial suspension. When indicated, PGF heat-inactivation may be performed (80°C, 10 min); the addition of a protease inhibitor cocktail may also be done following manufacturer instructions (Merck™ Complete™, Mini Protease Inhibitor Cocktail). Purified PGF was treated with hyaluronidase type I-S from bovine testes (ref H3506, Merck) for 1h at 37°C, when indicated.

**Purification of *Shigella* virulence factors and IcsA-containing membrane fractionation**

*SPATEs*. *E. coli* HB101 pSepA an *E. coli* HB101 pPic were grown overnight in LB medium with Ampicillin (100 µg.mL$^{-1}$) and subcultured in 500 mL fresh LB medium with Ampicillin (100 µg.mL$^{-1}$) at 37°C. Cultures were centrifuged for 20 minutes at 800 x *g* and the supernatant was filtered through a 0,22 µm diameter filter (TPP® Filtermax rapid bottle filter unit, ref. Z760900, Merck). Filtrated culture media were precipitated in 70% ammonium sulfate (ref. A4418, Merck). Precipitates were pelleted by centrifugation (7 500 x *g*, 30 min) and subsequently resuspended in 10 mL cold PBS. Samples were further dialysed in cold PBS and concentrated by centrifugation on 100kDa filter (Sartorius™ Vivaspin™ 20 Centrifugal Concentrator). Protein samples were stored at -20°C, as described previously.

*Ipas.* Purification of IpaB, IpaC and IpaD was achieved as described previously[3,15]. *S. flexneri* 5a was cultured at 37°C in 50 mL TSB culture medium until $OD_{600}$=0.4 was reached. The bacterial culture was centrifuged at 3.000 x *g* for 5 min and pelleted bacteria were washed twice in PBS and further resuspended in 200 µL PBS. Ipas secretion was induced by the addition of 0.05% congo red (ref. 6277, Merck). After 30 minutes at 37°C samples were centrifuged at 17.000 x *g* for 1 min. Ipas-containing supernatants were collected and stored at -20°C.

*IcsA-containing membranes.* An overnight culture of *S. flexneri* 5a was diluted at 1:100 in 200 mL of fresh TSB medium and incubated at 37°C until an $OD_{600}$ 0.4 was reached. The bacterial culture was centrifuged at 17.000 x *g* for 20 min and pellets were resuspended in 1mL PBS. Samples were sonicated (4 cycles of 30 seconds) and ultracentrifuged at 120.000 x *g* for 1h30 at 4°C. IcsA-containing pelleted membranes were resuspended in 1mL PBS and stored at -20°C.

**PGF proteolytic activity**

To assess PGF proteolytic activity, purified PGF (10µg) were incubated in PBS overnight at 37°C with 2 µg commercial purified proteins (lactoferrin (ref. L1294, Merck), IgA (ref. I4036, Merck), IgG (ref. I4506, Merck) and casein (ref. C3400, Merck)) or 10 µg *Shigella* bacterial virulence factors (SepA, Pic, IcsA and IpaB/C/D). Samples were separated by electrophoresis on 4-12% SDS-PAGE gradient gel (NuPAGE; Invitrogen). As indicated, proteins were stained with Coomassie (InstantBlue™ Coomassie Stain, Abcam) or analysed by western blot, as indicated. As a negative control, PGF proteolytic activity inactivation was achieved by heating (80°C, 10 min) or by the addition of a protease inhibitor cocktail (Merck™ Complete™, Mini Protease Inhibitor Cocktail) following manufacturer instructions.

**Microscopy**

*Light microscopy.* Bacteria (grown on agar pads) and neutrophil producing PGF (stained with Alcian blue or PAS) were imaged with Axioskop 2 light microscope (Zeiss, Germany) equipped with an Optikam Pro6 digital Camera (Optika, Italy), using a 100x (Plan-NEOFLUAR 100X 1.3NA Ph3 oil) and a 40x oil objective (Plan-NEOFLUAR 100X 1.3NA oil) respectively.

*Fluorescence microscopy*. Immunolabeled neutrophil secreting PGF and guinea pig tissues were imaged with a SP8 confocal microscope (Leica) using a 40x oil objective and the Leica Application Suite X (LAS X) software. Images were analysed with the Fiji software (ImageJ version 2.1.0/1.53j). Immunolabeled HEp-2 cells were imaged with an Axio Observer Z1 spinning disk (Zeiss) equipped with a CSU-X1 confocal scanner unit (Yokogawa). Images were acquired using a 40x Plan-apochromat oil objective 1.4NA (Zeiss) and were analysed with the Fiji software (ImageJ version 2.1.0/1.53j).

*Scanning Electron microscopy.* Neutrophil secreting PGF or releasing NETs were observed at 3,5 kV, in "high vacuum" mode, with a GeminiSEM 300 FESEM (ZEISS Germany).

**PGF vs NETs comparison by quantitative imaging analysis**

In order to make a distinction between PGF and NETs by a imaging quantitative analysis, fixed samples were stained with Myelotracker-Cy5 and DAPI and imaged with a confocal microscope. Both Myelotracker and DAPI signals were normalized using the 1st and 99.9th percentile as minimum and maximum limits. The cell body region of interest ($ROI_{cell}$) is computed using the Myelotracker channel. First, a Difference of Gaussian filter with corresponding sigma $\sigma 1=4$ and $\sigma 2=16$ for PGF and $\sigma 1=5$ and $\sigma 2=20$ for NETs. Then, a threshold using Otsu approach is applied, followed by a morphological binary dilation with a structural element of radius r=10 for PGF and r=2 for NETs. The region of interest is then used to remove any cell related intensity for both Myelotracker and DAPI channels. The remaining structure in the image, corresponding to the extracellular polymeric structures, is then detected by thresholding the Myelotracker channel minus the cell areas with the Otsu method. Both Myelotracker and DAPI intensities are collected for each pixel belonging to the polymeric structures. The distribution of Myelotracker and DAPI signal intensities was displayed for both PGF and NETs. A linear regression was used to compare the relationship between Myelotracker and DAPI signal intensities on both samples. A significant difference between PGF (0.0751 ± 0.009) and NETs (0.361 ± 0.009) was observed.

**Statistical analysis**

All statistical analyses have been performed with the Prism 9 software (GraphPad software, USA). Comparisons with two groups were supported with Student paired or unpaired-t tests, as indicated. When p-values for the significance of pairwise comparisons were calculated and displayed on figures, we used the following convention:

*ns not significant; $p \geq 0.05$;*$p < 0.05$ & $p \geq 0.01$; ** $p < 0.01$ & $p \geq 0.001$; *** $p < 0.001$*

**Data availability**

The data that support the findings of this study are available from the corresponding author upon request.

**References**

### Acknowledgements

A.C.A. was granted with a PhD fellowship from the University of Paris. We acknowledge support from the Biological Mass Spectrometry core facility at the University of Manchester. IBS acknowledges integration into the Interdisciplinary Research Institute of Grenoble (IRIG, CEA). This work was supported by the Agence Nationale de la Recherche ANR JCJC grant [grant number ANR-17-CE15-0012] and has benefitted from the support provided by the University for Advanced Study (USIAS) for a fellowship, within the French national program “Investment for the future” (IdEx-UNISTRA) (B.S.M.). This work was also supported by a grant from Medical Research Council [grant number MR/R002800/1] (D.J.T.). The Wellcome Trust Centre for Cell-Matrix Research, University of Manchester (D.J.T.), is supported by core funding from the Wellcome Trust [grant number 088785/Z/09/Z]. This work was also supported by the CNRS and the GDR GAG [GDR 3739], the “Investissements d’avenir” program Glyco@Alps [grant number ANR-15-IDEX-02], by grants from the Agence Nationale de la Recherche [grant number ANR-17-CE11-0040]. Electron microscopy imaging was performed on the Bordeaux Imaging Center, member of the FranceBioImaging national infrastructure, funded by the ANR [grant number ANR-10-INBS-04].

*In memory of Dr. Fabrizia Stavru (1978-2021).*

### Author contributions

B.S.M designed the study and supervised experiments. A.C.A performed the cell biology and biochemical experiments. M.S. performed the microbiology experiments. B.S.M. performed the animal experiments. R.V. performed GAG detection experiments. C.R and D.T. performed the mass spectrometry experiments. I.S.

performed the scanning electron microscopy experiments. S.R and J.-Y.T. performed quantitative imaging analysis. R.R.V. performed glycosaminoglycan analysis. P.J.S. analysed data and discussed findings. B.S.M. and A.C.A. wrote the manuscript. All authors edited and approved the manuscript.

### Competing interests

The authors declare no competing interests

### Additional informations

### Extended Data

#### Extended Data Fig. 1

**(a)** Neutrophils cultured for 4h under -$O_2$ conditions were fixed either with Carnoy's solution or PFA 3.3%. Samples were labeled with Myelotracker-Cy3 (red) and DAPI (blue) and imaged by confocal microscopy. Bars, 10 µm. **(b)** Kinetic study of PGF secretion by neutrophils (0, 4h and 18h, -$O_2$ conditions) in RPMI 1640 medium supplemented with 10 mM Hepes and 3 mM glucose. Samples were stained with alcian blue. Bars, 80 µm. **(c)** The viability of neutrophils secreting PGF (see (b)) was assessed by flow cytometry using propidium iodide (PI) staining. 'ns' indicates $p>0.05$ (Student's T-test, n=3) (**d**) As a control of Fig. 1e, non-infected guinea pig colonic mucosa was stained with DAPI (blue) and Myelotracker-Cy3 (red) and imaged by confocal microscopy. Bars, 10µm.

#### Extended Data figure 2.

**(a)** Extended list of proteins identified in purified PGF by mass spectrometry analysis, in complement to data presented in Fig. 2b. The subcellular localization of each protein is indicated ($\alpha$, $\beta$1, $\beta$2 or $\gamma$ granules) in addition to the number of unique peptide (1, 2-3 or >4) identified in each biological replicate (n=3) **(b)** PGF were separated on a SDS-PAGE gel (4-12%) or on an agarose gel (0.8%) and stained with Coomassie or transfer onto a PVDF membrane for western blot analysis, as indicated. The presence of myeloperoxidase, elastase, lactoferrin, cathelicidin (LL-37) and albumin in PGF was confirmed by western blot. **(c)** The accumulation of PGF most abundant proteins in vivo within *Shigella* foci of infection was confirmed by immunodetecting cathelicidin, elastase or myeloperoxidase (MPO) with specific antibodies (magenta) in the guinea pig colonic mucosa infected with *Shigella flexneri* 5a pGFP (green). Samples were additionally stained with DAPI (blue) and Myelotracker-Cy3 (red). Images were acquired by confocal microscopy. Bars, 20 μm. **(d)** The mRNA expression profiles of naïve neutrophils and neutrophils secreting PGF (18h culture under -$O_2$ conditions) were compared by RNAseq analysis (n=3). Results for *lactoferrin*, *lysozyme*, *cathelicidin*, *elastase*, *cathepsin G* and *myeloperoxidase* are highlighted in yellow.

**Extended Data figure 3.**

**(b)** The morphology of neutrophils secreting PGF (4h culture, -$O_2$ condition) and neutrophils releasing NETs (4h culture, +$O_2$ condition, +PMA) were compared by scanning electron microscopy (SEM) at x4.000, x10.000 and x25.000 magnifications. Bars, 1 μm. **(b)** The susceptibility of PGF to proteinase K treatment and not DNAseI was confirmed by immunofluorescence. Neutrophils were cultured for 4h in -$O_2$ condition and further treated with proteinase K or DNaseI for 1h at 37°C. Samples were fixed with a Carnoy's solution and labeled with DAPI (blue) and Myelotracker-Cy5 (magenta). Samples were imaged by confocal microscopy. Bars, 20μm.

**Extended Data Figure 4**

**(a)** *Shigella* infection induces neutrophil PGF secretion in +$O_2$ condition, as revealed by immunostaining. Similar results were obtained in -$O_2$ condition (see Fig. 3a). *Shigella flexneri* 5a was labelled with a specific α-LPS antibody (green), PGF was stained with Myelotracker-Cy3 (red) and DNA was stained with DAPI (blue). Bars, 50 μm. **(b)** The purified PGF proteolytic activity was abolished in the presence of a cocktail of protease inhibitors or upon heat-inactivation (control of data presented in Fig. 3c). Treated PGF was incubated for 1h at 37°C with SepA or Pic, which were separated on

SDS-Page gel and stained with Coomassie. **(c)** The stability of human lactoferrin and immunoglobulins (IgG and IgA heavy/light chains) in the presence of PGF was confirmed after 1h co-incubation at 37°C. Proteins were separated on SDS-Page gel (4-12%) and stained with Coomassie.

**Extended Data Figure 5**

**(a)** *E. coli* K12, *Shigella flexneri* 2a and *Shigella sonnei* growth inhibition by purified PGF was demonstrated by culturing bacteria in the presence of $O_2$ on agar pads in the presence of PGF (37°C, 4h and 18h). Bars, 5 µm. The confluence of bacterial cultures was quantified using the Fiji software (n=3, * indicates $p < 0.05$, ** indicates $p < 0.01$ and *** indicates $p < 0.001$, Student's T-test). **(b)** The inhibition of *Shigella flexneri* 5a growth by PGF is comparable in the presence or in the absence of $O_2$. Bacteria were cultured with PGF on agar pads (37°C, 4h and 18h). Bars, 5 µm. The confluence of bacterial cultures was quantified using the Fiji software (n=3, * indicates $p < 0.05$, and *** indicates $p < 0.001$, Student's T-test).

**Extended Data Figure 6**

**(a)** The cytotoxic activity of purified PGF was assessed by incubating human epithelial HEp-2 cells with purified PGF for 6h or 24h. Low [PGF] corresponds to 30 µg purified PGF/mL and High [PGF] corresponds to 300 µg purified PGF/mL. The impact of PGF on Hep-2 cell morphology was assessed by immunofluorescence. Treated cells were fixed in PFA 3.3%. Actin was stained with phalloidin-FITC (green) and DNA was stained with DAPI (blue). **(b)** The adherence of treated cell was assessed by quantifying the confluence of cell cultures using the Fiji software (n=3, *** indicates $p < 0.001$, Student's T-test). **(c)** The viability of Hep-2 cells upon PGF treatment was assessed by flow cytometry using a PI staining. Cells were treated as indicated in (a) and were trypsinized prior analysis. **(d)** % of cell death corresponds to the proportion of PI-positive cells (n=3, ** indicates $p < 0.01$, Student's T-test).

**Extended Data Figure 7**

**(a)** The presence of chondroitin sulfate or keratan sulfate in purified PGF was analyzed by western blot using specific antibodies, after electrophoresis on O.8% agarose gel or 4-12% SDS Page gel. **(b)** The presence of chondroitin/chondroitin sulfate in NETs was confirmed by RPIP-HPLC. Heparan sulfate, chondroitin di- or tri-sulfate were not detected. Results are expressed as percentage of total NETs GAGs (n=3).
