## Extended Figures for "Neutrophils degranulate GAG-containing proteoglycofili, which block *Shigella* growth and degrade virulence factors"

### Extended Data Fig.1

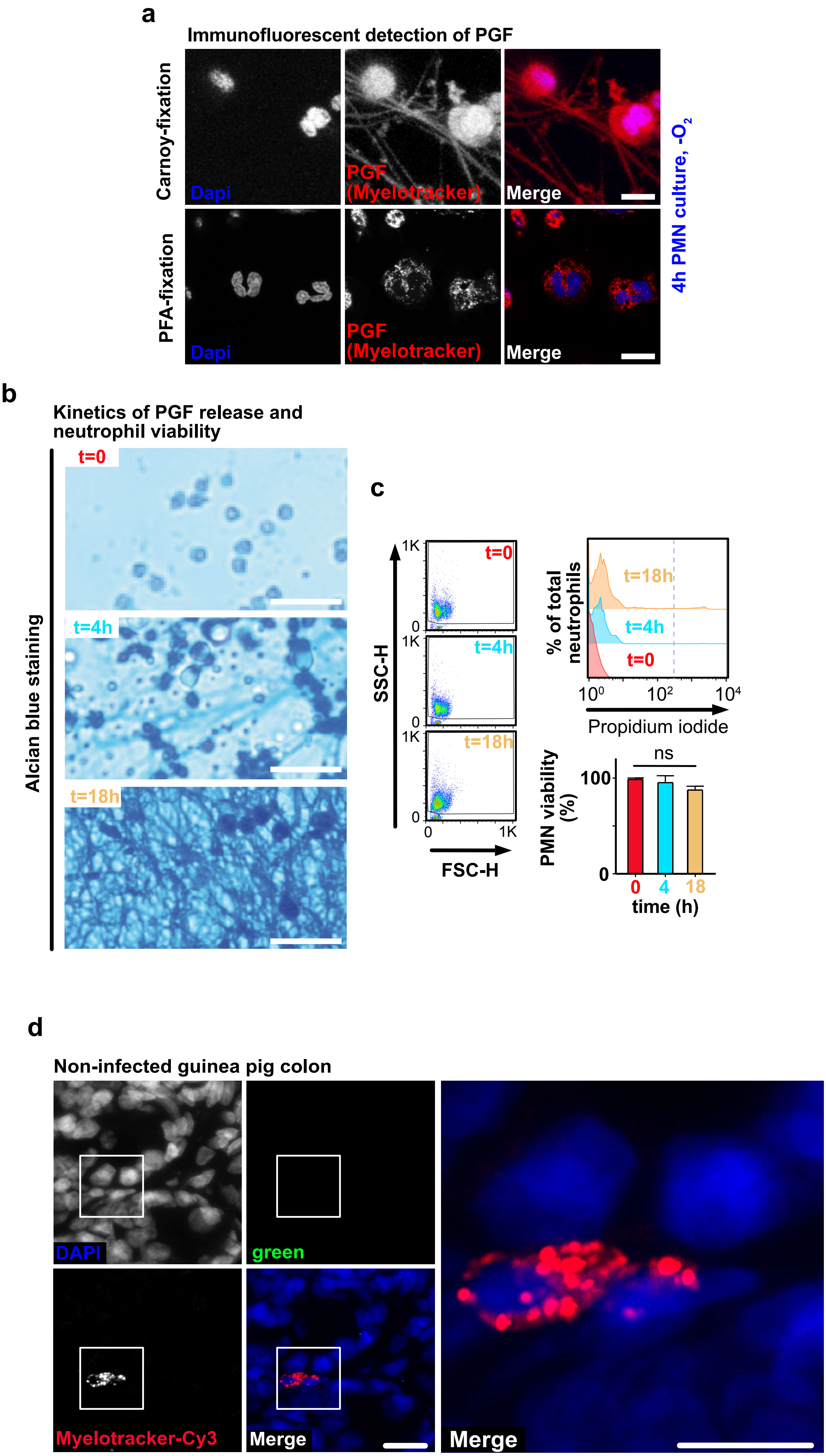

#### Extended Data Fig.2

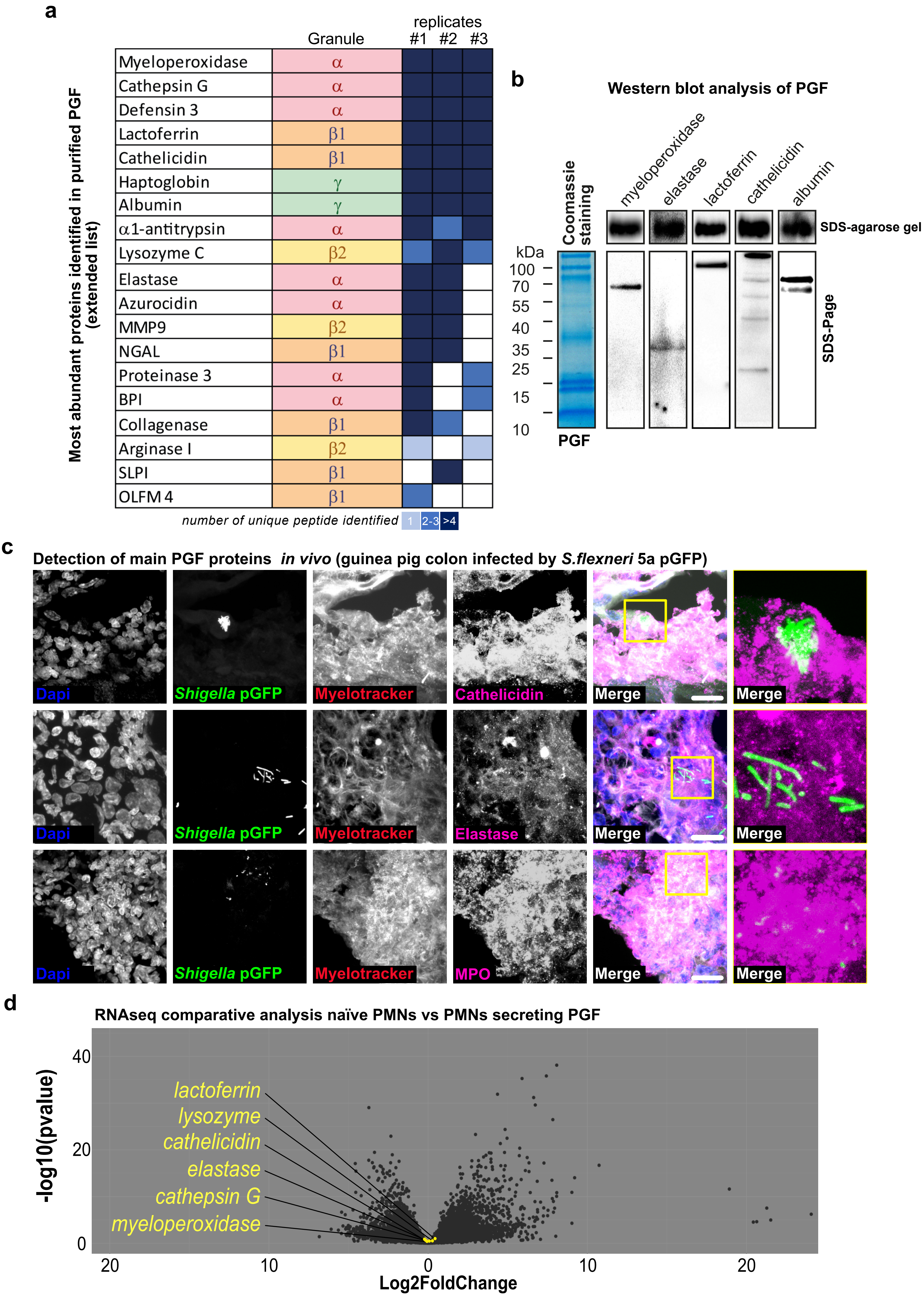

### Extended Data Fig.3

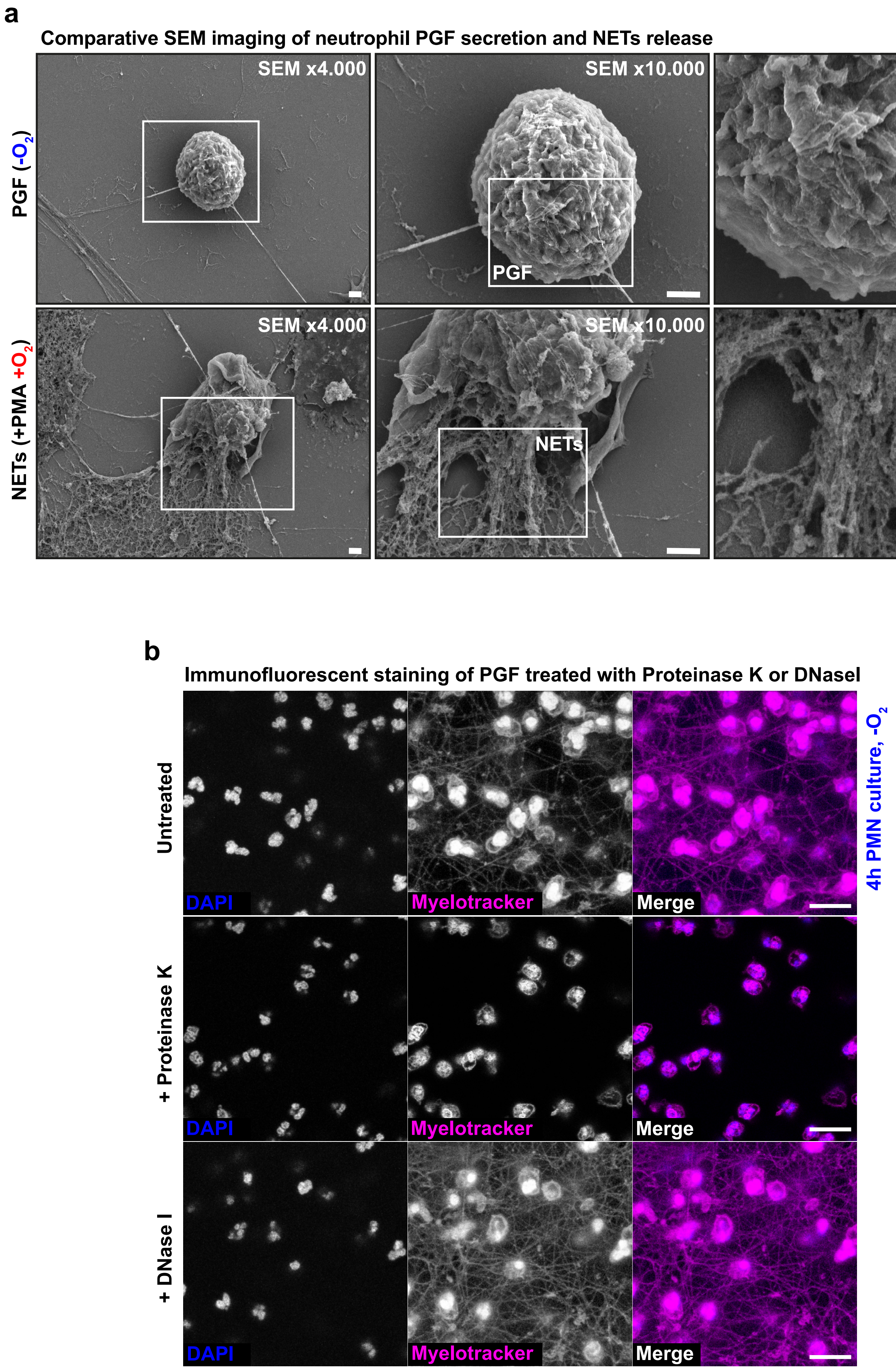

#### Extended Data Fig.4

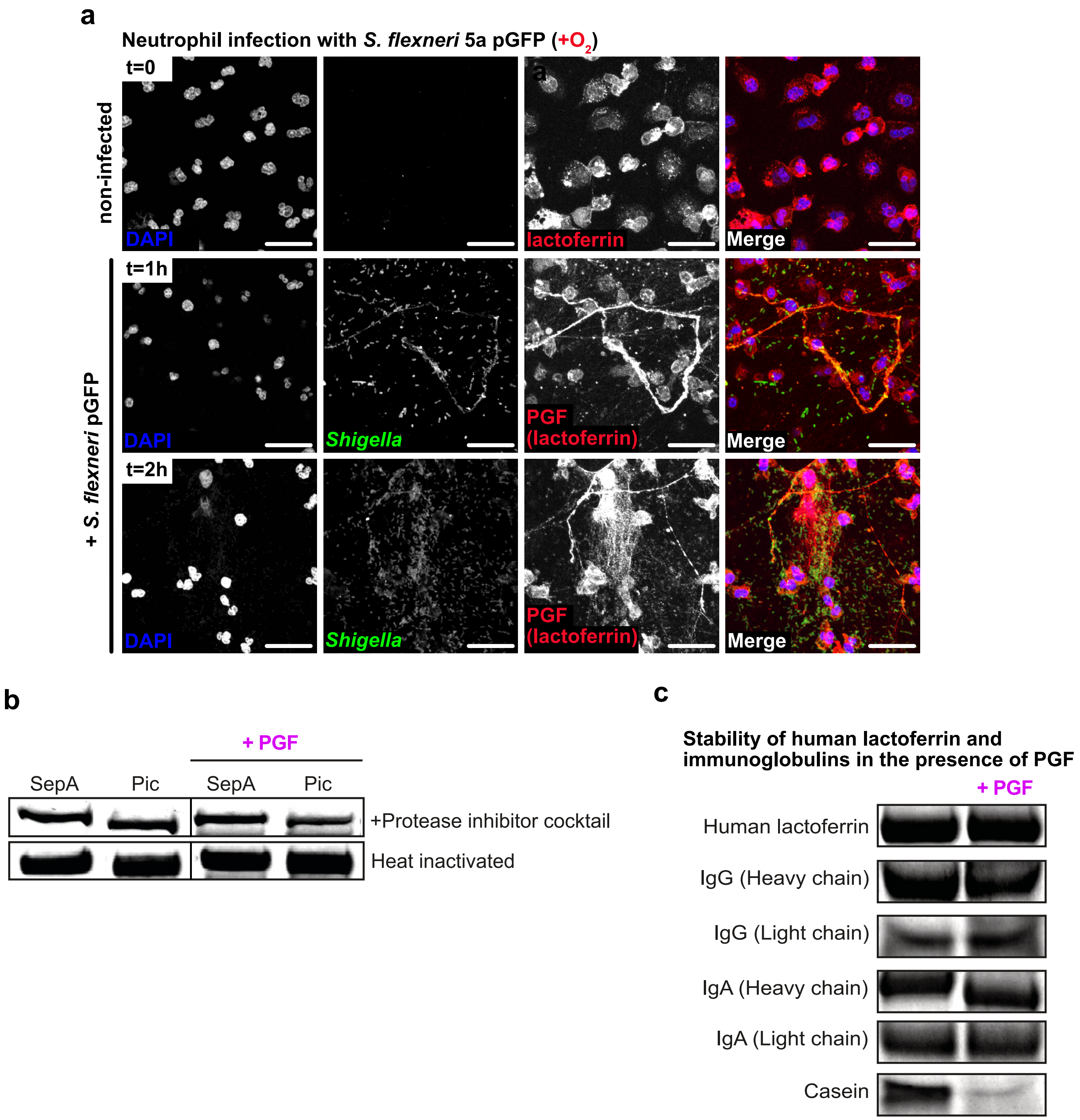

### Extended Data Fig.5

a

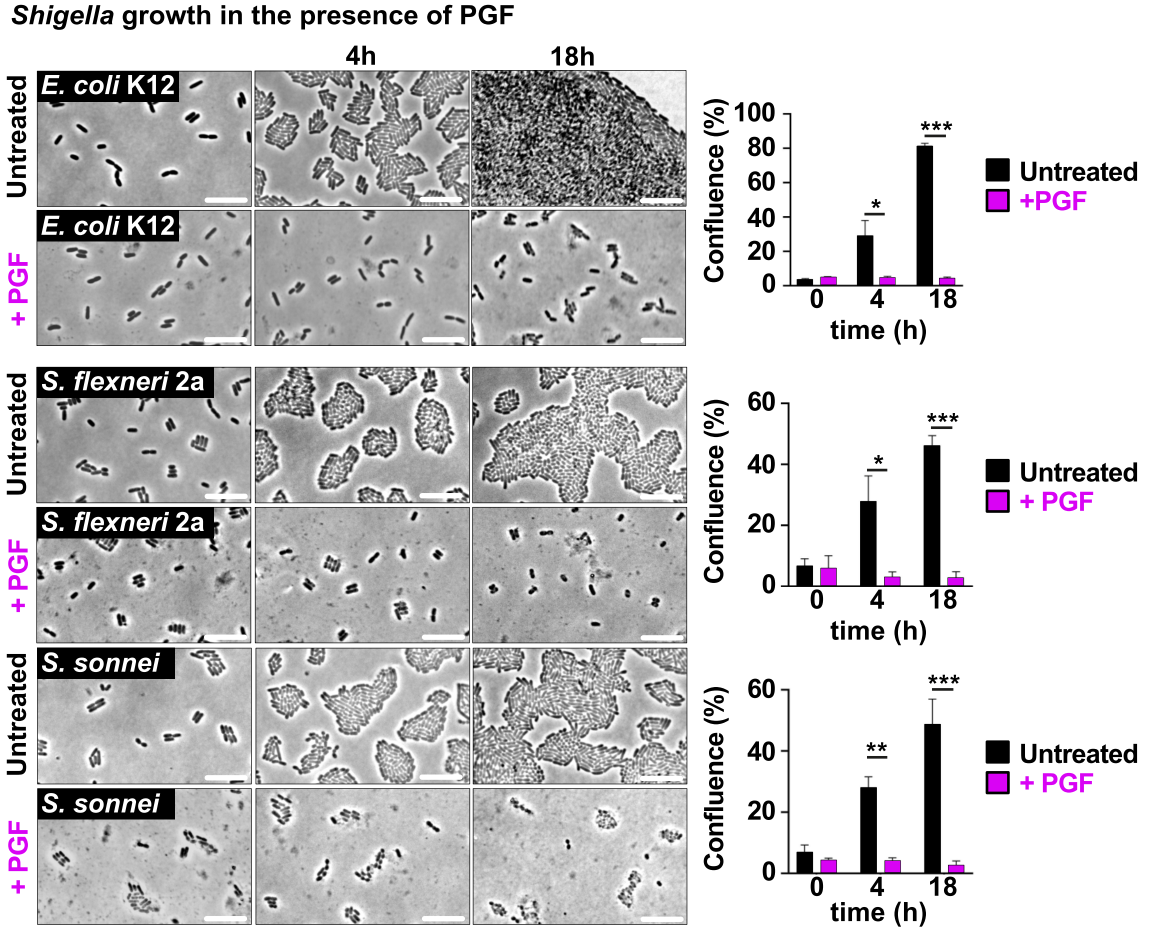


b

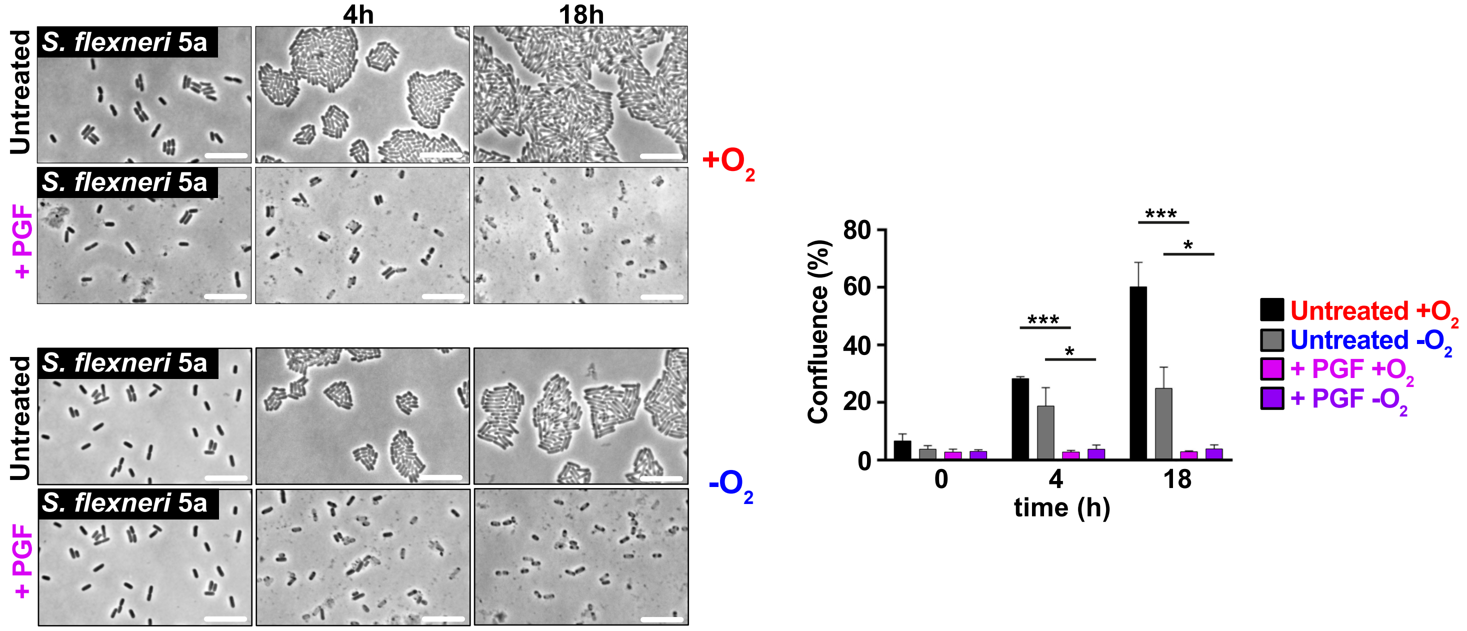

**Extended Data Fig.6**

**a**

**Immunofluorescence analysis of HEp-2 cells treated with PGF**

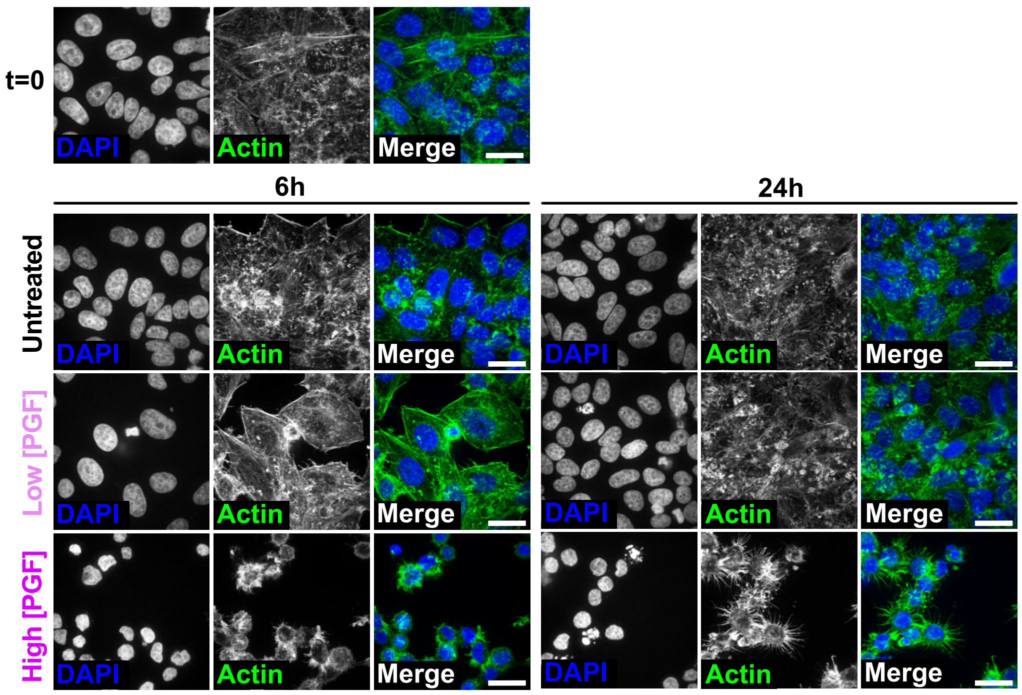


**b**

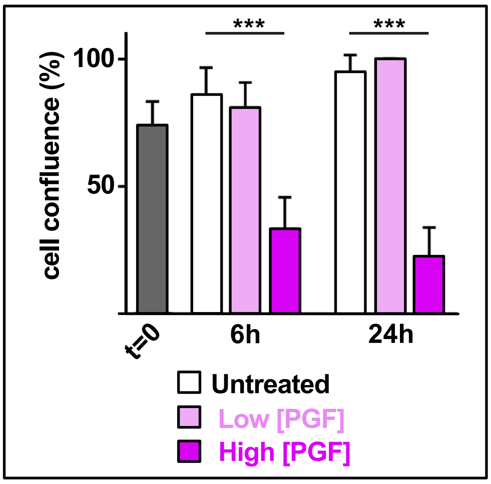


**c**

**Flow cytometry analysis of HEp-2 cells treated with PGF**

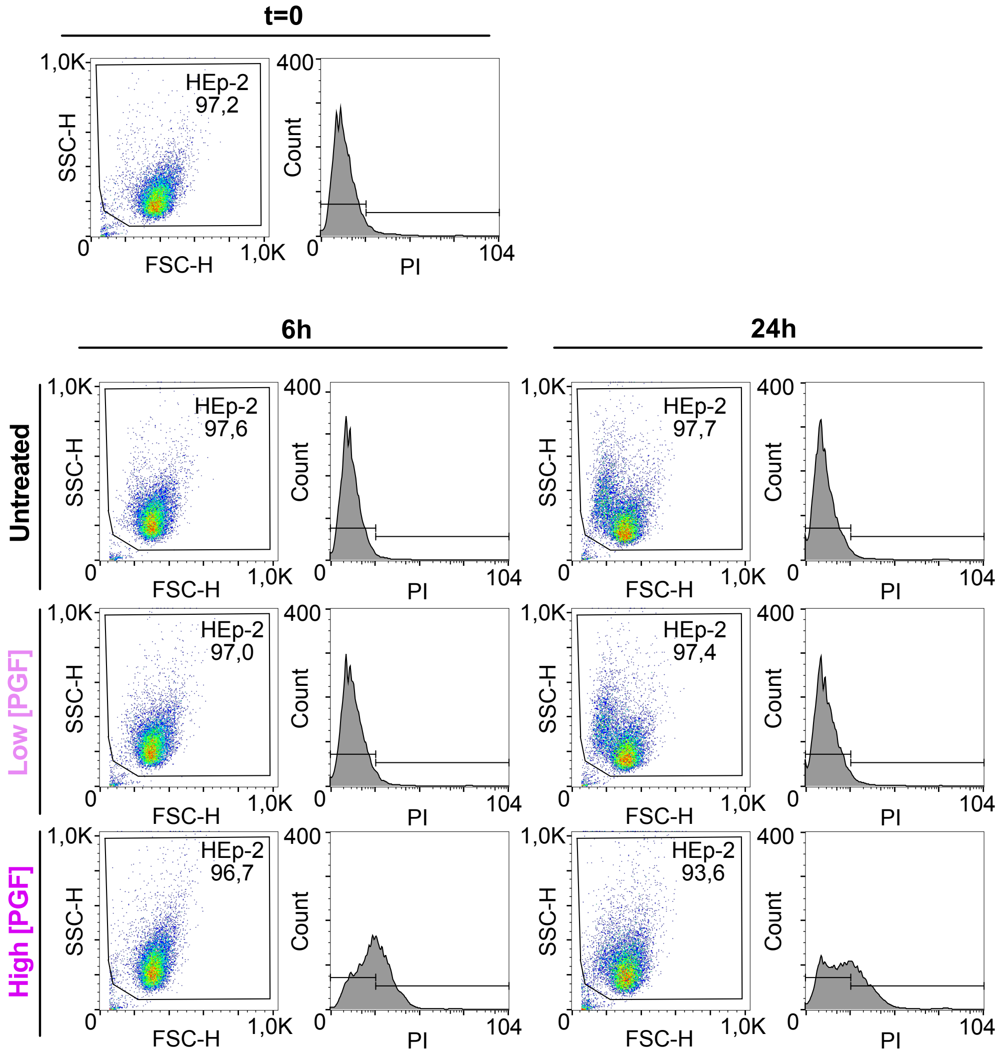


**d**

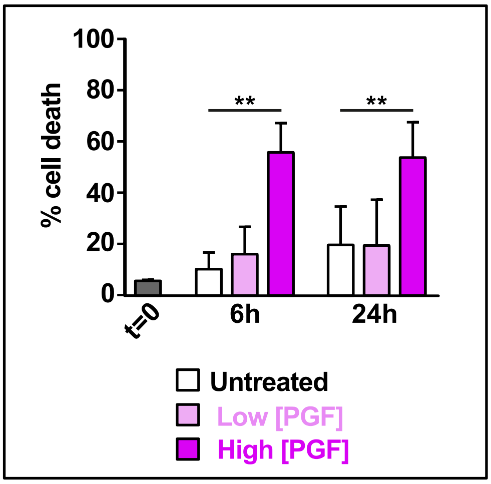

#### Extended Data Fig.8

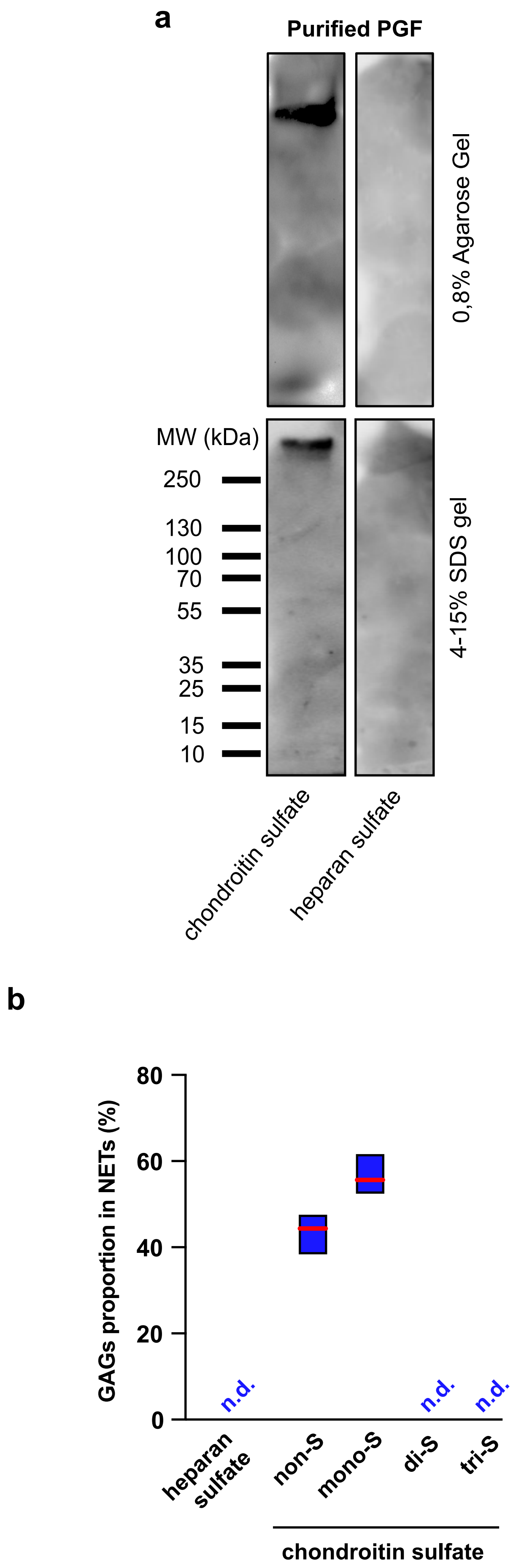
